# The study of fibrotic scar at the long term spinal cord lesion rats

**DOI:** 10.1101/2023.01.03.522540

**Authors:** Maryam Naghynajadfard, Ying Li

## Abstract

The fibrotic lesion formation is the main deterrent at regeneration of neurons in CNS injuries. In this paper we studied fibrotic scar formation in rat corticospinal tract lesion after one year survival time. The glial scar formation and Extracellular Matrix (ECM) fibronection derived scaffolds were investigated and the fore paw reaching task was performed during this period to see whether there is any regeneration of neurons along the lesion or not.

## Introduction

Although Fibrotic scar is necessary to wound healing to tissue injury, it is followed by loss of function and damaged neurons not being replaced by regeneration in brain and spinal cord lesion (Bradbury EJ., et al., 2019) and (Jun JI., et al., 2018). Fibrotic scar also calls mesenchymal scar contains components such as Fibroblast/ Fibroblast-like cells, Extracellular Matrix (ECM) including Collagen I, IV, Fibronectin, Laminins and others such as EphB2, Phosphacan, NG2, Tenascin and Semaphorin III (Ayazi M., et al., 2022). Fibrotic scar is limited by glial scar which contains high level of astrocytes which is worked as an impediment for exonal regeneration. Little is known about the formation of acute and chronic lesion in rat CNS. However, as the studies in mice shows fibroblasts accumulate in the lesion center following five days after injury and starts increasing by day 7 which is followed by the rise of machrophage in the lesion area which triggers ECM protein accumulation in lesion (Soderblom, C., et al., 2013) (Zhu, Y., et al., 2015). The matured fibrotic scar is formed n CNS under trauma by 14 days post-injury and the scar persist chronically 56 days post injury(Zhu, Y. et al., 2015) (Komuta, Y., at al., 2010). Acute injury can lead to wound repair with tissue replacement when in chronic injuries it can lead to overtime increasing tissue alteration (Burda JE. et al., 2014). Understanding the difference in chronic and acute fibrotic scar can help with solving many problems. As this article we will look at the fibrotic scar at corticospinal tract lesion in rats after one year post surgery to achieve more understanding about the mechanism of fibrotic scar at long term injury.

## Material method

In our study animal were used in accordance with the UK Home Office regulations for the care and use of laboratory animal, the Uk Animals (Scientific Prosecures) Act 1989, with the ethical approval of the University College London, Institute of Neurology.

5 adult female rats (180-210 g body weight) of a locally inbred Albino Swiss Strain (AS) were under unilateral corticospinal cord lesion using KCTE-TC-S electrode with a straight RF tip and kept as control for the period of one year to study fibrotic scar. The directed forepaw reaching (DFR) was counted once a week. Animal were perfused and tissue were prepared and cut coronally and horizontally. Immunohistochemistry staining was performed. Poly clonal rabbit anti-human Fibronectin (Dako, Uk), was used to stain fibronectin and anti-Glial Fibrillary Acidic protein clone GA5 (Chemicon, Uk) was used to study astrocyte behaviour.

## Result

The animals have been under spinal cord surgery to produce corticospinal tract lesion in order to cause deficite at paw reaching task. The lesion has destroyed Cu: Cuneate fascuculusm and gr: Gracile fasciculus Figure 1(B&C). The animals lost paw reaching task and tested once a week for one year. The result showed that animal did not have any return in paw reaching and remained the deficiency.

**Fig 1:**
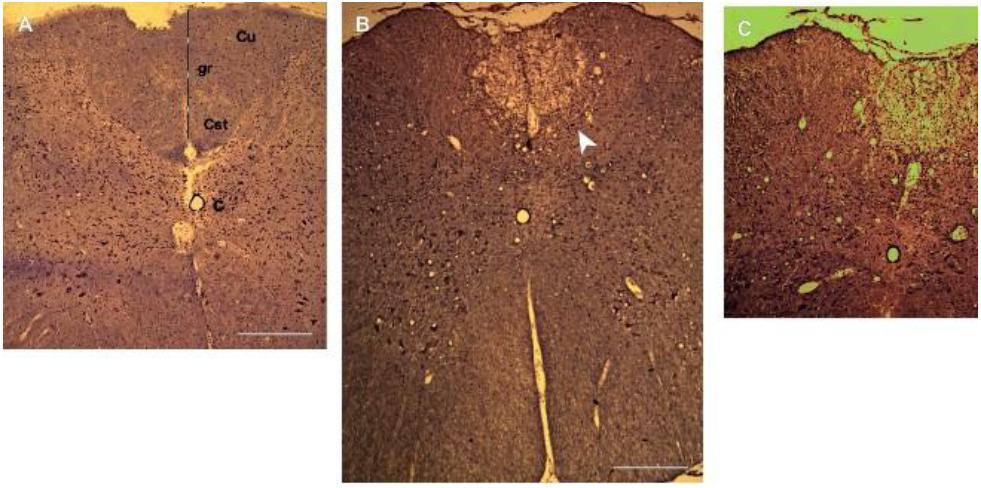
A: 20-μm-thick coronal section of long term lesion Cu: Cuneate fascuculusm, gr: Gracile fasciculus 400μm B: corticospinal tract lesion 200μm C: Cortic lesion, 100 μm

The animals being killed after one year and tissue prepared for cross sectioning. The sectioned being labelled by fibronection (Red) to visualized Fibrotic scar and the asctrocyets have been labelled by Green GFAP to present glial scar figure2 (A,B&C).The glial scar bordered fibrotic scar and fibronection have been scattered at lesion core.

**Fig 2.**
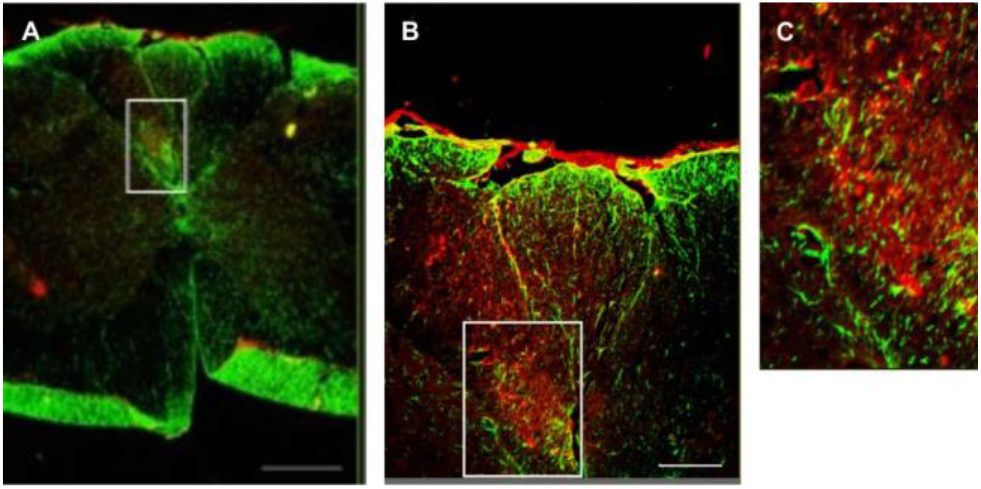
A. 20-μm-thick coronal section of long term lesion GFAP (Green) and anti-fibronectin (Red) positive (arrows). (B and C) the enlarged view of lesion area, intense GFAP immunoreactivity around the lesion leading to a dense, and “closed” scar completely walling off the central lesion area with astrocytic process passing through the fibrotic scar. C. hypertrophic fibronectin response in lesion central, Survival time: 8 months. Scale bar; 500μm; 200 μm; 50 μm.

The animals have been under transplant of Olfactory ensheating cells (OEC) 4 months after destruction of constricspinal tract. The results showed that animals regenerate axons at the margin of lesion Figure 3 (A) and spread around glial scar (Green A&B). That proves that regeneration of axons can occur along glial scar by transplanting OECs.

**Fig 3:.**
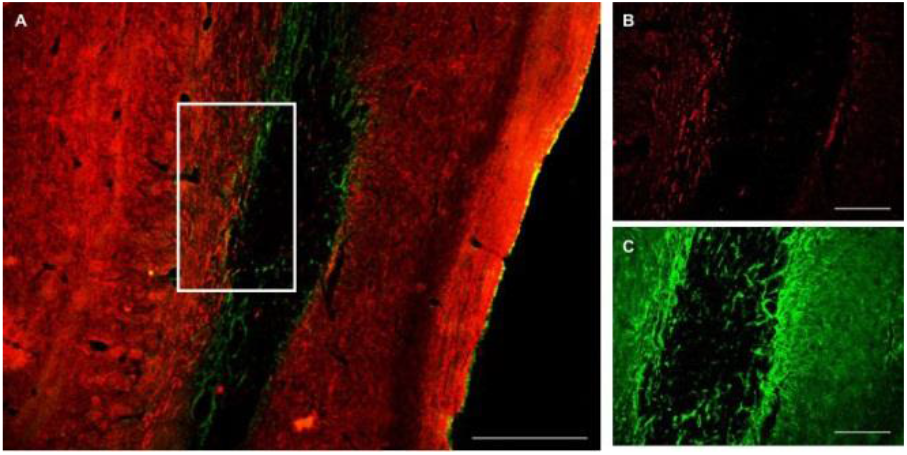
A. Reconstructed corticospinal tract after transplantation of OECs, (axons NF, Red). B & C. Enlargement of regenerated NF labelled axons (red, outlined in A) and Astrocytes, Glial Scar Horizontal section, Survival time: 4 months, Scale bar: 200 μm; 100 μm; 100 μm.

## Discussion

After spinal cord injury (SCI) tissue is going through healing process which leads to formation of fibrotic scar, glial scar and deposition of ECM which limits the regeneration of axon due to creation of inhibitory molecules and producing a physical barrier that avoids axonal regeneration in between the lesion core (Dias, D.O. et sl., 2018) (Fernandez-Klett, F., et al., 2014) (Dias, D.O., et al., 2018). Fibrotic scar majorly is made from components such as Colagen, Fibronection and Laminine (Zhu et al., 2015) (Yokota et al., 2017). In rat SCI causes cavity formation in lesion core which indeed speared in smaller area than mice (Buss et al., 2007) (Zhu et al., 2015). In rat fibrotic scar exists along the edge of the cavity and partly join with astrocytes scar which may prove that astrocytes are involved in creation of fibrotic scar in rat (Zhu et al., 2015). When asctrocytes activate they deposite ECm compntent, Chrondrotin sulphate proteoglycans (CSPGs) which creates glial scar that districts fibrotic scar after SCI (Anderson et al., 2016). In GFAP-STAT3-CKO mice in which STAT3 is dysfunction, the hypertrophy of asctrocytes is missed and it caused the destruction of astrocytic scar which results in distruption of boundery with fibrotic scar. This indicate the crosstalk between fibrotic scar and glial scar after SCI (Renault-Mihara et al., 2017) (Dias et al., 2018). After Transplant of Olfactry ensheating cells in rat survival time of 4 months costicospinal tract lesion, OECs modulates the lesion environment and remodel reactive asctrocytests leading to regeneration of asctrocyts at the edge of lesion area. In deed glial scar acts as a bridge to permit regeneration axons pass through lesion area (Guijarro-Belmar et al., 2019). Another strategy is that the phenotype of asctrocyes has changed (Hara et al., 2017). For example, at animal model of neurodegenerative disease the disruption of neuron and behaviour loss has improved by blocking microglia-mediated A1 astrocytic alteration (Yun et al., 2018). This is one of the possibilities of conversion after transplant of OECs.

## Funding

The study was supported by the British Neurological Research Trust and the international Spinal Research Trust.

## Declaration of competing interest

The authors declare that they have no known competing financial interest or personal relationships that could have appeared to influence the work reported in this paper.

## Notes

### Competing Interest Statement

The authors have declared no competing interest.

